# Development and Characterization of Co-crystals Assisted with *In-silico* Screening for Solubility and Permeability Enhancement of Curcumin

**DOI:** 10.1101/2023.09.08.556876

**Authors:** Meera Singh, Shweta Takawale, Rajesh Patil, Somdatta Chaudhari, Aarifkhan Pathan, Jaiprakash Sangshetti, Abu Tayab Moin, Talha Zubair, Mohammad Helal Uddin

**Author notes:** For Correspondence: Mohammad Helal Uddin Professor Department of Applied Chemistry and Chemical Engineering (ACCE), Faculty of Science, University of Chittagong, Chattogram- 4331, Bangladesh.

## Abstract

Despite being a promising phytochemical, Curcumin’s potential applications are limited due to its classification in BCS class IV, which is associated with low water solubility and permeability. Enhancing the bioavailability of BCS class IV drugs presents a significant challenge, but crystal chemistry provides a hopeful avenue for overcoming this hurdle. In this research, co-crystals of Curcumin were developed to improve both solubility and permeability. Unlike traditional methods that require extensive trial-based lab work and time-consuming screening of co-formers, the use of molecular docking in *In-silico* co-former screening offers a scientific and rational approach to selecting suitable partners. In this study, two distinct co-crystals were synthesized using a solvent evaporation technique with methanol as the solvent, employing a 1:1 molar ratio. L-proline and piperine were chosen as co-formers to enhance solubility and permeability, respectively. The co-crystals underwent optimization and characterization through Design of Experiments (DOE). Comparing the dissolution study results for the same curcumin concentration, the cumulative drug release (CDR) after 8 hours was 20% for pure curcumin and an impressive 71% for curcumin-L-proline co-crystals. The permeability study, conducted over four hours using the everted gut sac method in phosphate buffer pH 6.8, revealed curcumin’s permeability to be less than 0.05 mg/mL, while curcumin-piperine co-crystals exhibited a five-fold increase (0.2545 mg/mL) in permeability. The co-crystals formed through a molecular ratio of 1:1 for curcumin-L-proline to enhance solubility and 1:1 for curcumin-piperine to enhance permeability, both demonstrated positive outcomes with support from optimization analysis, FTIR, DSC, SEM, PXRD analysis, and dissolution studies.

## 1. Introduction

The use of poorly soluble drugs has limitations because of low and irregular bioavailability, which leads to decreased safety and efficacy, particularly when administered orally[1,2]. Approaches to address these challenges have thus been thoroughly researched and can be divided into two categories: structural engineering (e.g., salt or prodrug formation) and formulation strategies (e.g., particle size reduction, addition of surfactant, co-solvation, amorphous solid dispersion or lipid-based formulation etc.) [3–9] Amongst them, salt formation is one of the most well-established ways for altering the aqueous solubility of an active pharmaceutical ingredient (API), but it has a drawback that it is applicable to only ionizable molecules [10]. Co-crystallization is also a promising approach for altering the water solubility of API when salt formation is not possible [11–18]. Supramolecular chemistry is the origin of co-crystallization, which occurs when two or more molecules are present in a unit cell which consists of non-covalent interactions like hydrogen bonds, pi–pi interactions and van der waals interactions between molecules. Pharmaceutical co-crystals, unlike other techniques, can be made from non-ionizable medicines without affecting their pharmacological properties. For medications with a low solubility, pharmaceutical co-crystallization is an effective technique to improve solubility and dissolution profile [19–23]. Co-former with higher hydrophilicity is able to increase the solvent affinity of crystalline phase, which results in better solubility of the poorly water soluble drug. Beside solubility, permeability is another key physicochemical feature that impacts oral absorption and helps the drug maintain a favorable pharmacokinetic profile. Several theories about how co-crystals help a drug overcome its poor permeability profile have been proposed by various research groups [24]. USFDA defines co-crystals as "crystalline materials composed of two or more molecules, one of which is the API and other is co-former, in a defined stoichiometric ratio within the same crystal lattice that are associated by nonionic and non-covalent bonds [25,26]." Choosing the correct co-formers for each drug is one of the most difficult aspects of creating pharmaceutical co-crystals. There have been several methods for selecting co-formers and screening co-crystals, such as Hydrogen-bonding propensity, Synthonic engineering, Supramolecular compatibility by Cambridge Structure Database (CSD), pKa based models, Fabins’s Method, Lattice energy calculation, Hansen solubility parameter, Thermal analysis measuring saturation temperature etc. but each method has its own set of limitations. Stability in the presence of excipients is also a hurdle in the development of co-crystals. Molecular Docking is one method for virtual screening of co-formers. The ability to form reversible or non-covalent interactions with the API is used to choose co-formers. Hydrogen bond donors (HBDs) or acceptors (HBAs) should be present in both the API and the co-former, such as ether, thio-ether, alcohol, ketone, thio-ketone, ester, thioester, carboxylic acid, amide, primary and secondary amines, and so on. Van der Waals forces, pi-pi (-) interactions, aromatic stacking, and halogen bonding are some of the other interactions involved in the production of co-crystals. Molecular docking helps the formulator to understand the type of intermolecular interaction between drug and co-former by using docking and virtual analysis leading to selection of best suitable co-former in less time and resources [19]. Thus, molecular docking is a method for fast screening of more than one co-formers at a time and also understanding the type of intermolecular interaction of drug and co-former. This method of screening also helps to locate the position of the hydrogen bond formed and displays its binding energy along with its distance. It gives a better understanding to the formulator to predict the chemical structure of co-crystals in the 3- dimensional view.

## 2. Results and Discussion

### 2.1. Molecular Docking study

A negative bond energy between the drug and co-former indicates the potential for co-crystal formation. Hydrogen bonding exhibits greater strength than Pi-Pi bonding. The software indicates a positive interaction energy when co-crystal formation is unlikely [27]. AutoDock software analysis revealed that among the 28 co-formers tested, piperine exhibited a binding energy of -3.68, while L-proline displayed a binding energy of -2.52 (**Table 1**). Notably, both values were lower compared to the other co-formers. Piperine demonstrated favorable pi-pi and pi-alkyl interactions, whereas L-proline formed one hydrogen bond with curcumin. Based on these findings, piperine was chosen as the co-former to enhance permeability, while L-proline was selected to improve solubility.

**Table 1.**
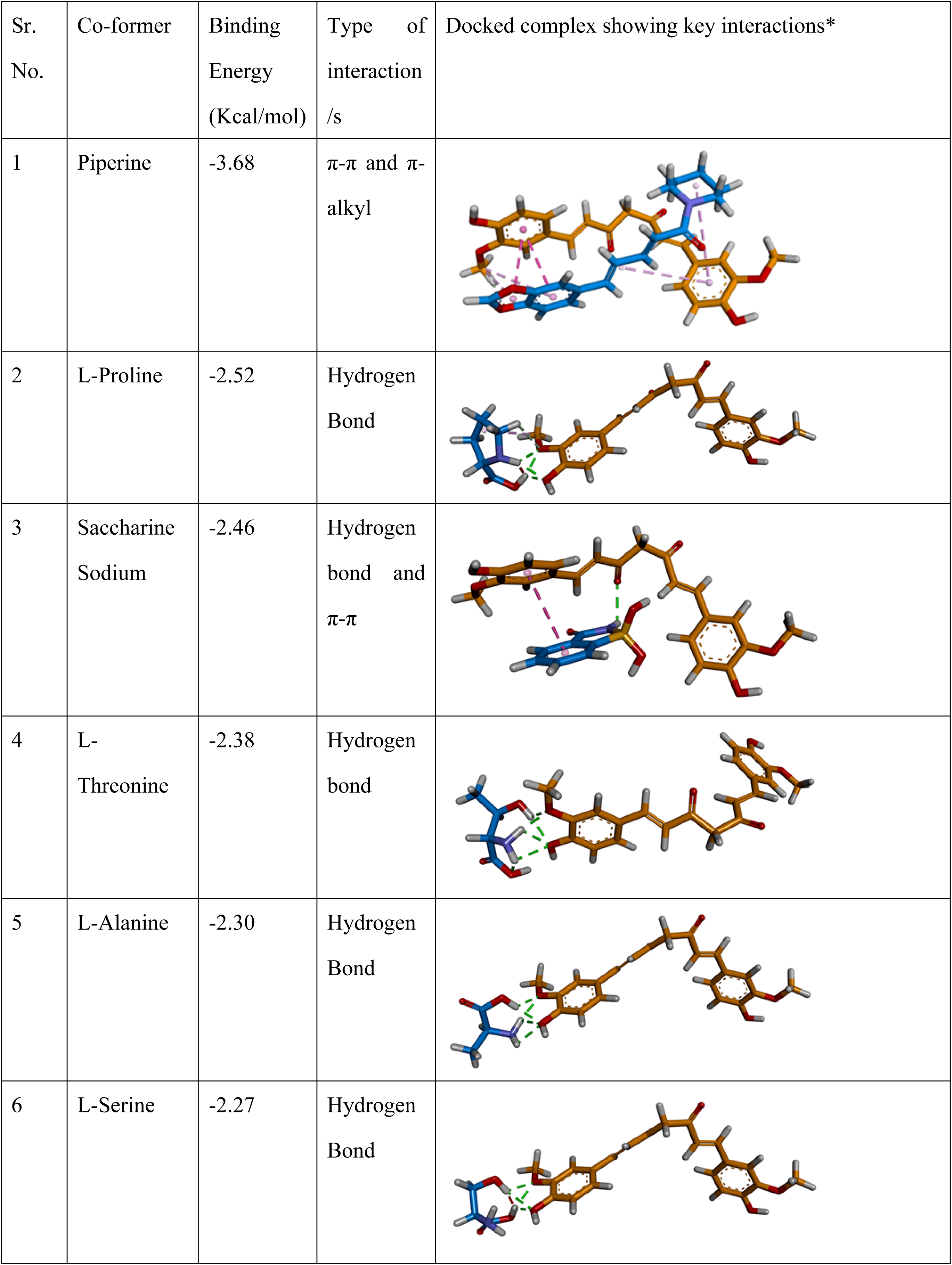
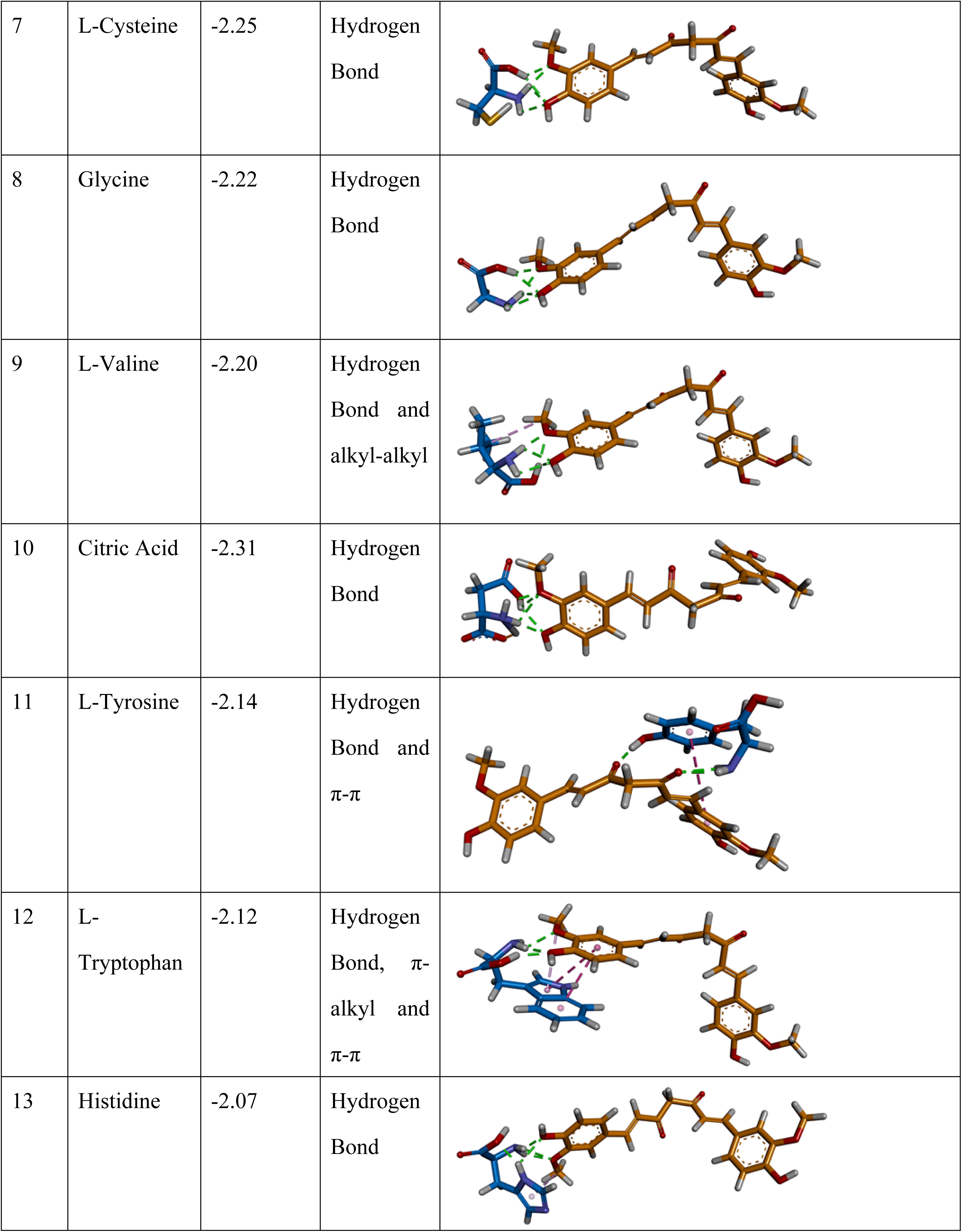
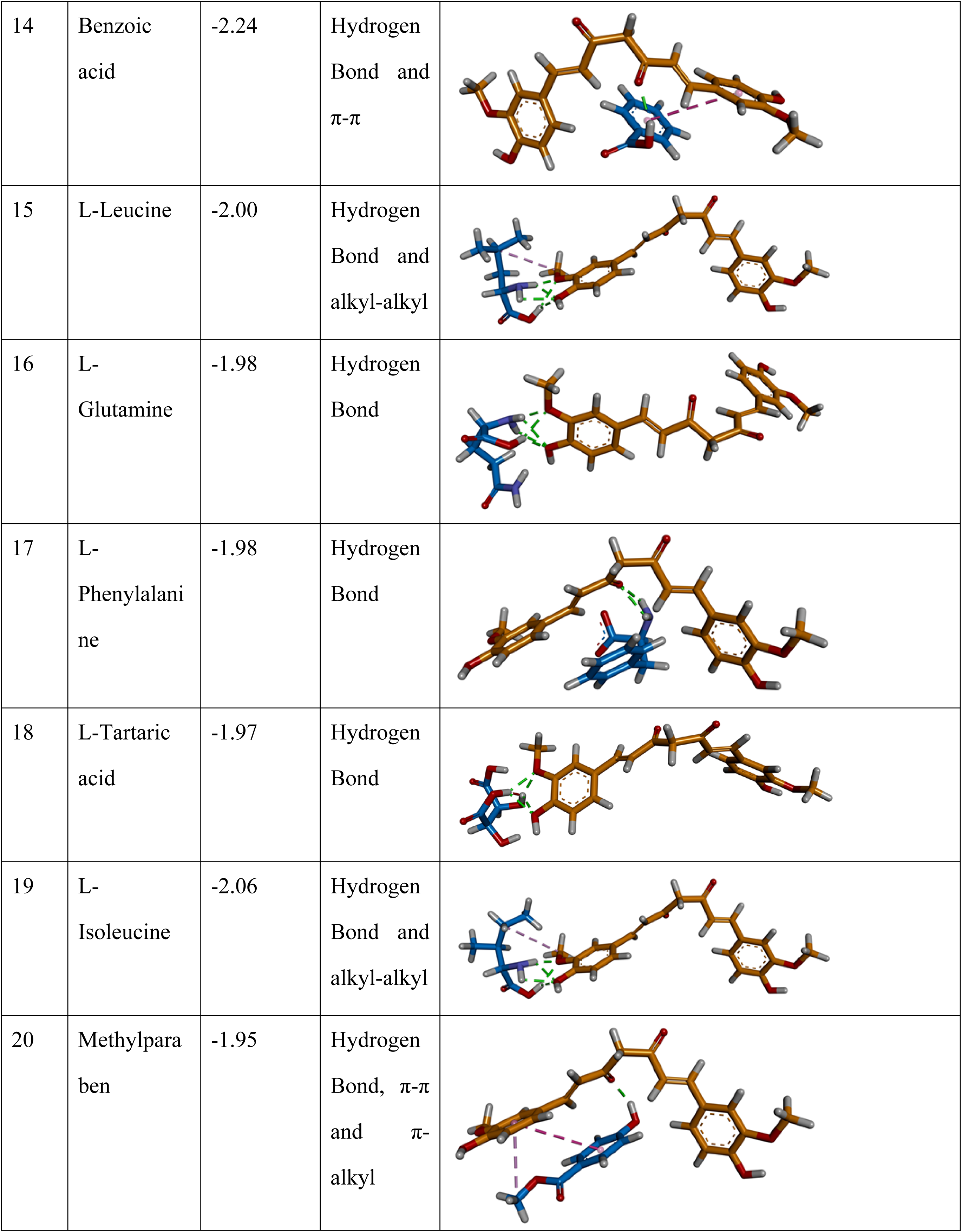
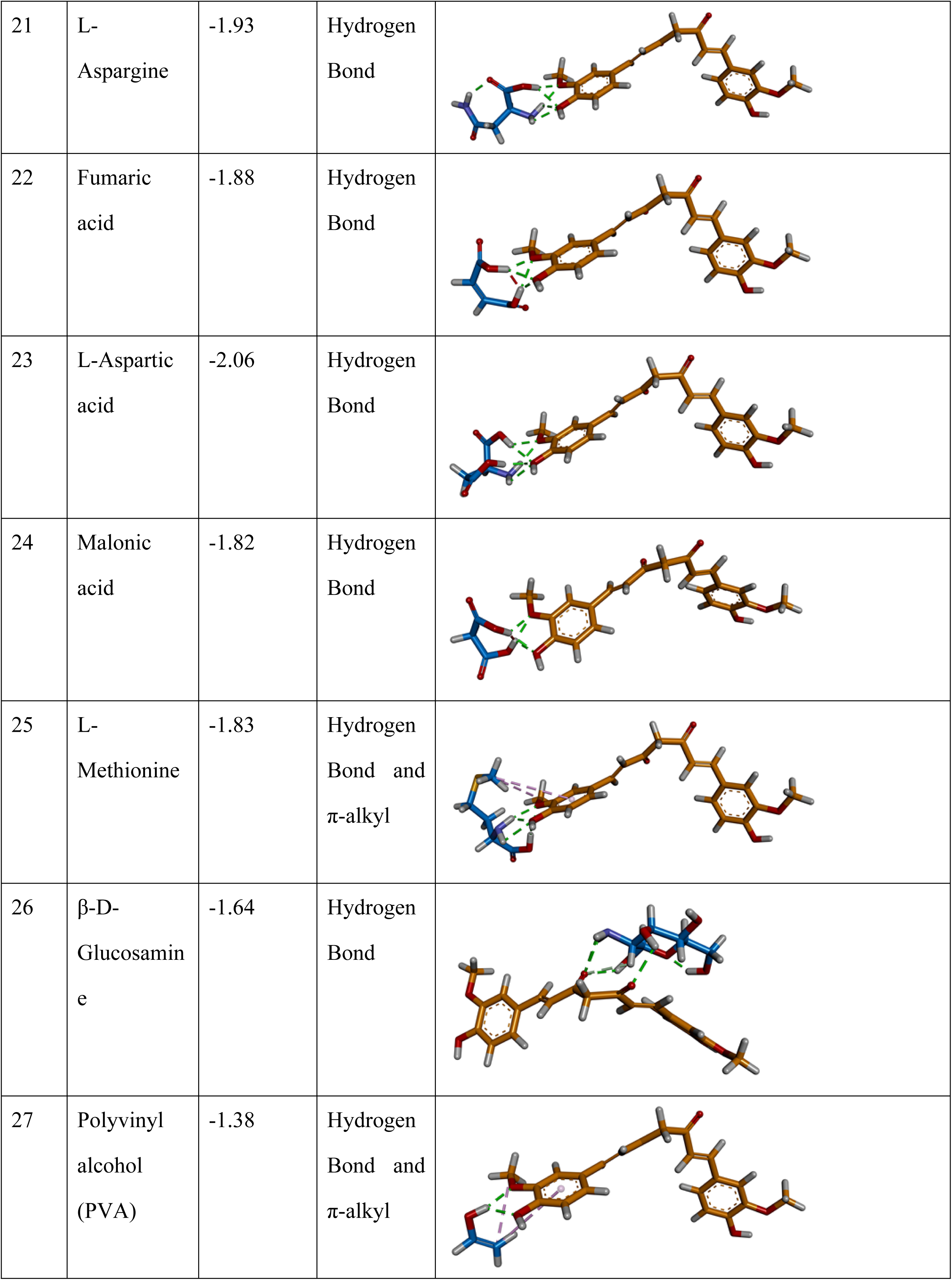
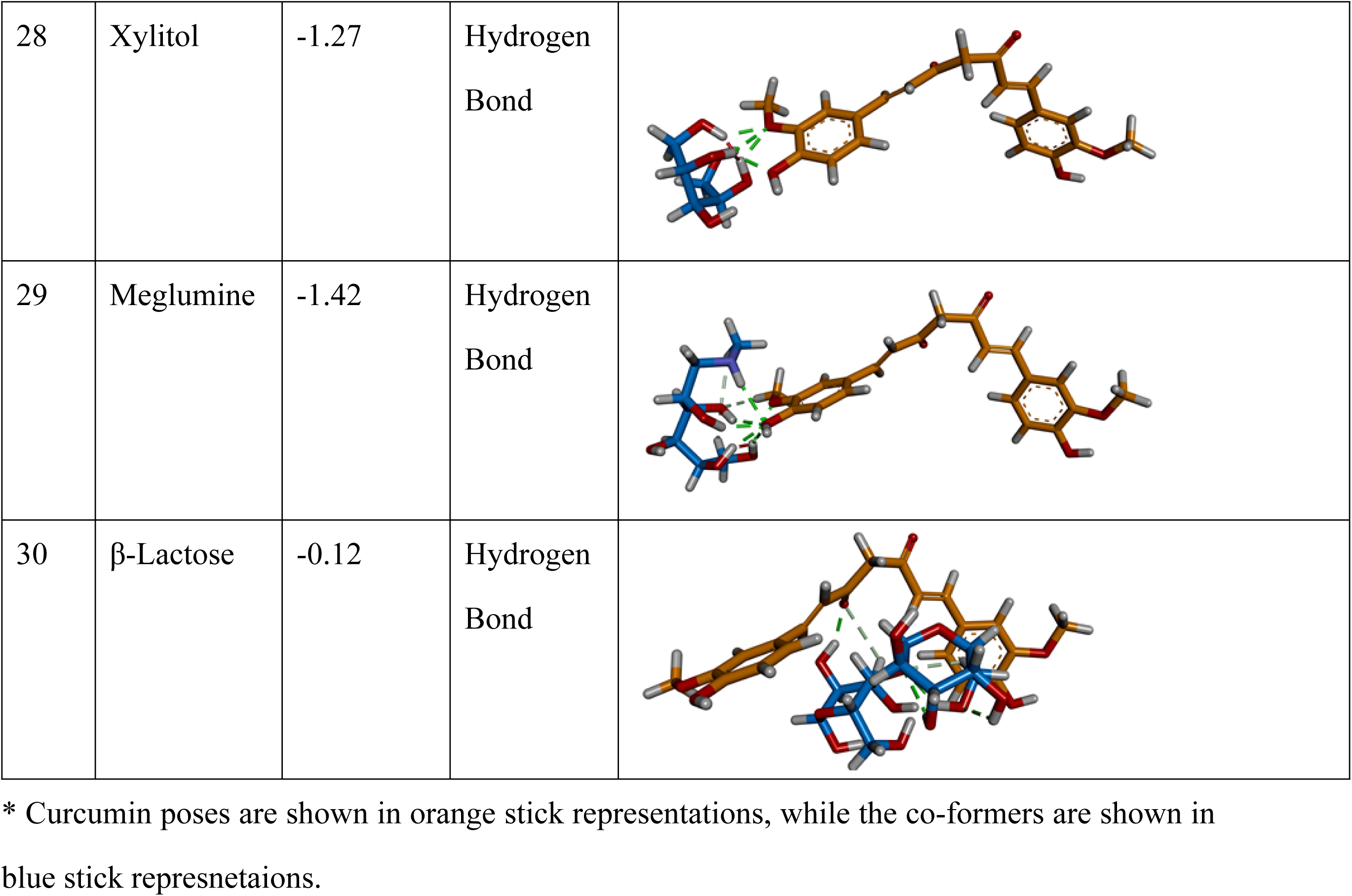
Outcomes of *In-silico* Molecular Docking showing promising co-formers.

| Sr. No. | Co-former | Binding Energy (Kcal/mol) | Type of interaction /s | Docked complex showing key interactions* |
| --- | --- | --- | --- | --- |
| 1 | Piperine | -3.68 | π-π and π-alkyl | |
| 2 | L-Proline | -2.52 | Hydrogen Bond |  |
| 3 | Saccharine Sodium | -2.46 | Hydrogen bond and π-π | |
| 4 | L-Threonine | -2.38 | Hydrogen bond |  |
| 5 | L-Alanine | -2.30 | Hydrogen Bond |  |
| 6 | L-Serine | -2.27 | Hydrogen Bond |  |
| 7 | L-Cysteine | -2.25 | Hydrogen Bond |  |
| 8 | Glycine | -2.22 | Hydrogen Bond |  |
| 9 | L-Valine | -2.20 | Hydrogen Bond and alkyl-alkyl |  |
| 10 | Citric Acid | -2.31 | Hydrogen Bond |  |
| 11 | L-Tyrosine | -2.14 | Hydrogen Bond and π-π | |
| 12 | L-Tryptophan | -2.12 | Hydrogen Bond, π-alkyl and π-π | |
| 13 | Histidine | -2.07 | Hydrogen Bond |  |
| 14 | Benzoic acid | -2.24 | Hydrogen Bond and π-π | |
| 15 | L-Leucine | -2.00 | Hydrogen Bond and alkyl-alkyl |  |
| 16 | L-Glutamine | -1.98 | Hydrogen Bond |  |
| 17 | L-Phenylalanine | -1.98 | Hydrogen Bond |  |
| 18 | L-Tartaric acid | -1.97 | Hydrogen Bond |  |
| 19 | L-Isoleucine | -2.06 | Hydrogen Bond and alkyl-alkyl |  |
| 20 | Methylparaben | -1.95 | Hydrogen Bond, π-π and π-alkyl | |
| 21 | L-Asparagine | -1.93 | Hydrogen Bond |  |
| 22 | Fumaric acid | -1.88 | Hydrogen Bond |  |
| 23 | L-Aspartic acid | -2.06 | Hydrogen Bond |  |
| 24 | Malonic acid | -1.82 | Hydrogen Bond |  |
| 25 | L-Methionine | -1.83 | Hydrogen Bond and π-alkyl | |
| 26 | β-D-Glucosamine | -1.64 | Hydrogen Bond | |
| 27 | Polyvinyl alcohol (PVA) | -1.38 | Hydrogen Bond and π-alkyl | |
| 28 | Xylitol | -1.27 | Hydrogen Bond |  |
| 29 | Meglumine | -1.42 | Hydrogen Bond |  |
| 30 | β-Lactose | -0.12 | Hydrogen Bond | |

\* Curcumin poses are shown in orange stick representations, while the co-formers are shown in blue stick representations.

### 2.2. Application of DOE after preparation of preliminary batches

Two separate 3^2^ factorial designs used to find out the correct ratios of coformers for solubility enhancement and permeability enhancement showed B3 and B4 batches as optimized batches with Lproline and piperine as coformers respectively. These batches are considered opimized because of following DOE results.

### 2.3. *In Vitro* Dissolution Rate Profiles of suggested nine batches by DOE

The dissolution rate of the co-crystals was significantly enhanced, as depicted in **Figure 1A and 1B**. In **Figure 1A**, the co-crystals (batch B3), formulated in a molar ratio of 1:1 [100:31.2 mg], exhibited the most substantial dissolution rate improvement when compared to both pure curcumin and the other 8 co-crystal batches. Over an 8-hour period, batch B3 demonstrated a curcumin dissolution percentage of 71%, whereas only 20% of pure curcumin dissolved within the same duration. This outcome was further validated and subjected to analysis utilizing Design of Experiments (DoE) to confirm the optimization of batch B3.

**Figure 1.**
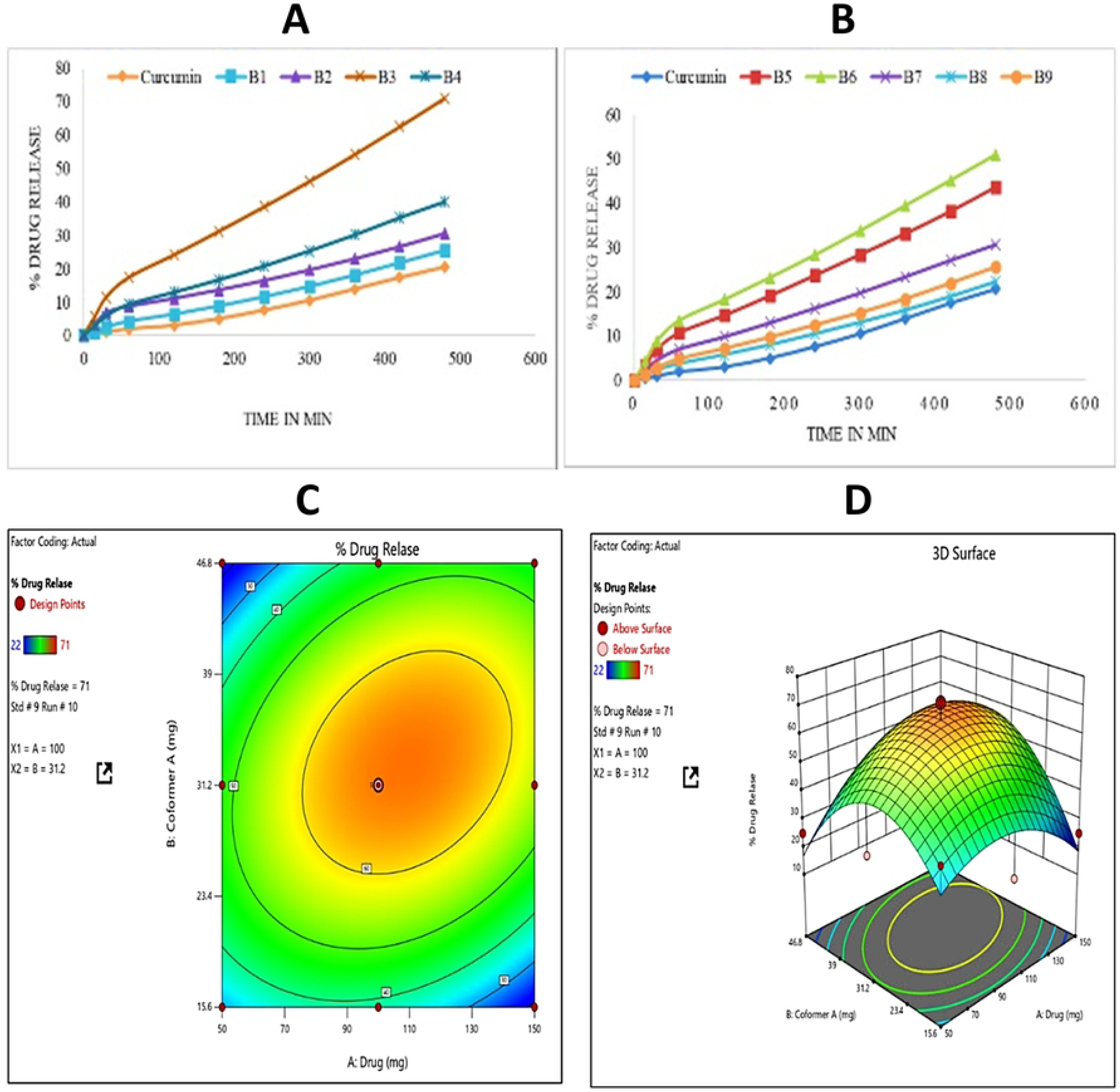
(A) In-vitro dissolution Rate Profile of curcumin and B1, B2, B3and B4 (B) In-vitro dissolution rate profile of curcumin and B5, B6, B7, B8, B9 (C) 2D and (D) 3D Surface Graph for % Drug Release

### 2.4. ANOVA for Quadratic model [Response: % drug release]

The prepared co-crystals were divided into 9 batches. The analysis of these batches revealed that the model exhibited an F-value of 4.09, indicating its significance. The probability of an F-value of this magnitude occurring due to random variations is only 4.09%. Model terms are considered significant when their corresponding p-values are less than 0.0500. The fit summary suggests a quadratic model. Upon comparing predicted and actual % CDR (cumulative drug release) values, it is evident that the software’s predictions closely align with the experimental values obtained from the dissolution study. In **Figure 1C and 1D**, the 2D and 3D representations of % drug release are depicted. The optimized batch, identified as std.4 - run 10, features a curcumin quantity of 100 mg and L-proline quantity of 31.2 (with a molar ratio of 1:1), resulting in a % drug release of 71%.

The final ANOVA equation, marked as equation (1), is as follows:

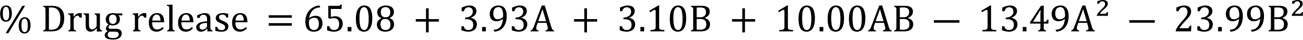

ANOVA for quadratic model for Response: % drug release showed that the quadratic model is significant with P value of 0.0467.

The quadratic model has better values for suggested and adjusted R^2^ than other models.

### 2.5. Ex-vivo Permeability Study using Everted Gut Sac Method

The permeability of curcumin in the co-crystal form with piperine exhibited a significant improvement. In **Figure 2A and 2B**, the co-crystal batch with piperine (B4) at a molar ratio of 1:1 [100:77mg] demonstrated the highest permeability compared to pure curcumin (0.0142 mg/mL) and the other 8 batches. Approximately 0.2545 mg/mL of curcumin concentration permeated from batch B4 co-crystals after 4 hours. **Figures 2C and 2D** depict the 2D and 3D representations of the percentage of drug permeation.

**Figure 2.**
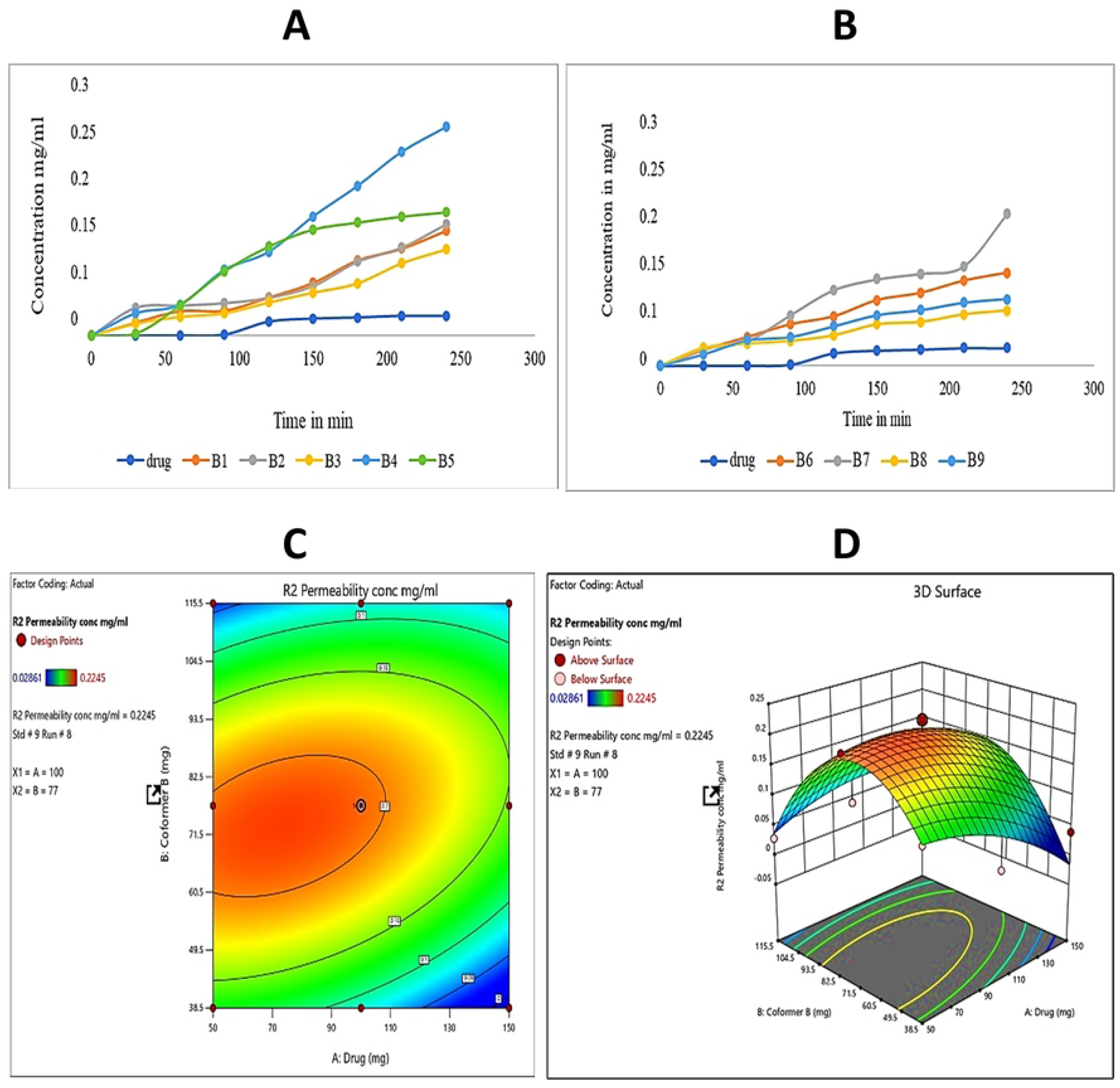
(A) Ex-vivo permeability of curcumin and B1,B2,B3, B4 and B5, (B)Ex-vivo permeability of curcumin and B6, B7, B8 and B9 (C) 2D and (D) 3D Surface Graph for permeability mg/mL

### 2.6. ANOVA for Quadratic model [Response: Permeability in concentration mg/mL]

The Model F-value of 4.45 indicates the significance of the model. Model terms are considered significant when their p-values are less than 0.0500.The fit summary suggests a Quadratic model, as displayed.

In **Figure 2C and 2D**, both 2D and 3D representations depict the amount of permeated concentration in mg/mL. Among the various batches, run 4 with a standard of 5 was optimized. This batch contained 100 mg of curcumin and 77 mg of piperine, with a molar ratio of 1:1. The permeated concentration was measured at 0.2545 mg/mL.

The final ANOVA equation, labeled as equation (2), for permeability in concentration is as follows:

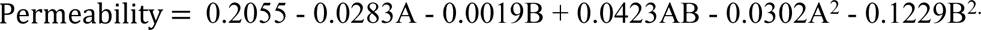

ANOVA for quadratic model for Response: permeability showed that the quadratic model is significant with P value of 0.0383. The quadratic model has better values for suggested and adjusted R^2^ than other models.

### 2.7. Saturation Solubility Study

The saturation solubility of curcumin and curcumin-L-proline co-crystals was assessed and juxtaposed with the documented solubility of curcumin in water. The results revealed that the solubility of curcumin in distilled water was 0.07928 mg/mL, closely resembling the reported solubility (**Table 2**). In contrast, a fourfold surge in solubility was noted in the co-crystal configuration, reaching 0.47654 mg/mL.

**Table 2.** Comparative solubility study of curcumin and curcumin co-crystals.

| Sr. No. | Compound | Solubility<br>(mg/ml) | Reported solubility<br>(mg/ml) |
| --- | --- | --- | --- |
| 1 | Curcumin | 0.07928 | <0.1 |
| 2 | Curcumin –L-proline co-crystals | 0.47654 | - |

### 2.8. Fourier Transform-Infrared Spectroscopy Analysis-curcumin, L-proline and cocrystals

The peaks observed at 3749.74 cm−1 and 1026 cm−1 are indicative of O-H stretching resulting from hydrogen bonding between an oxygen atom of curcumin and the hydrogen atom of L-proline. The shifted frequencies strongly imply their participation in robust intermolecular hydrogen bonding within the co-crystal phase. More comprehensive information is available in **Table 3 and Figure 3A**.

**Figure 3.**
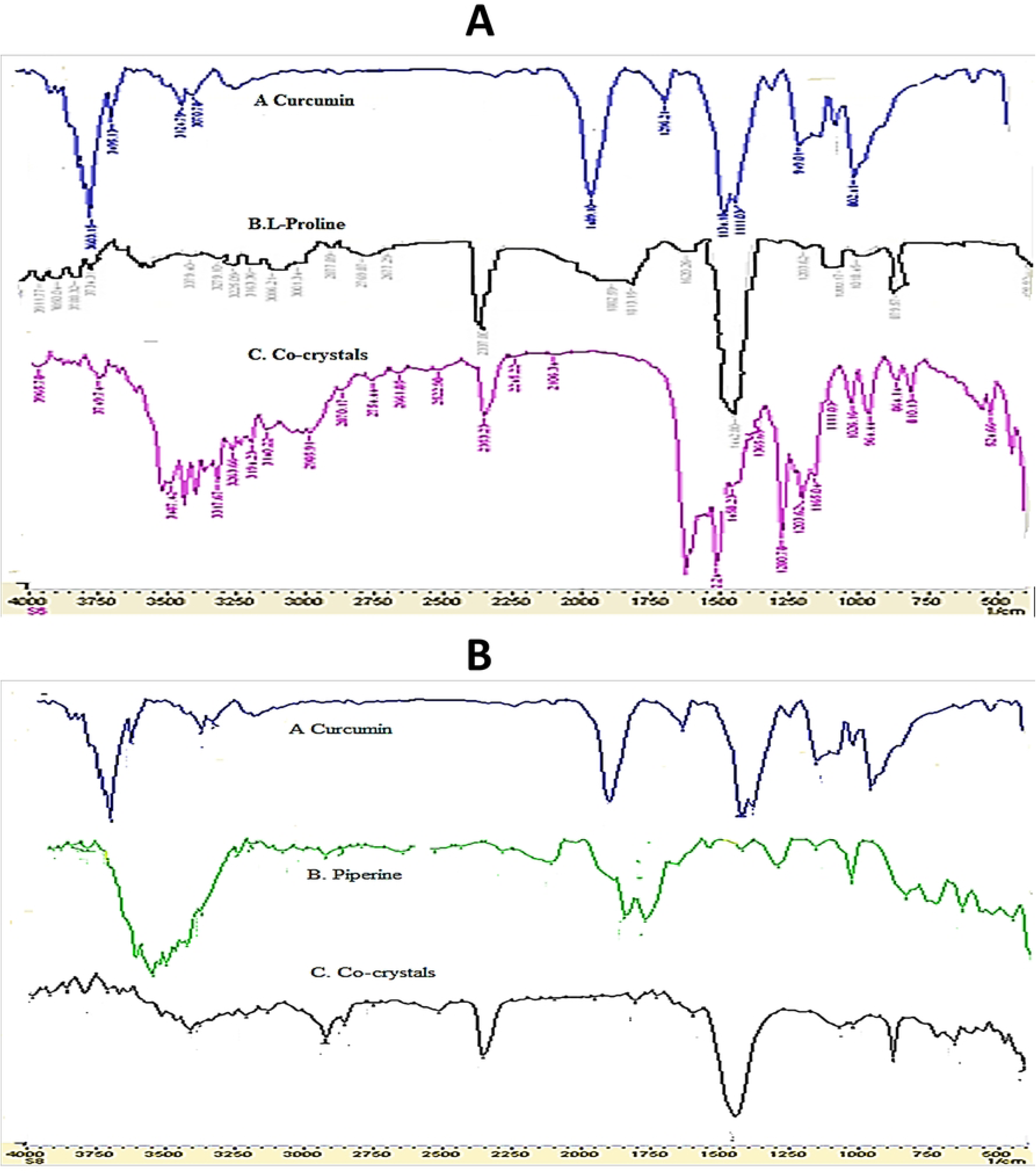
Fourier Transform-Infrared Spectra of Curcumin, L-proline, and their respective co-crystals: (A) Curcumin-L-proline and (B) Curcumin-Piperine

**Table 3.** Observations of Fourier Transform-infrared spectra of (A) Curcumin(B) L-proline, and (C) the cocrystal of curcumin-L-proline.

| Sr. No. | Sample | Bond | Wavenumber (cm-1) |
| --- | --- | --- | --- |
| 1 | Curcumin | OH Stretching | 3603 |
|  |  | Aromatic (C=O and C=C) stretching | 1509 |
|  |  | C-H stretching | 1489 |
|  |  | Aromatic C-O | 1296 |
|  |  | C-O-C stretching | 949 |
| 2 | L-Proline | OH stretching | 3734 |
|  |  | Aliphatic C-H stretching | 2337.80 |
|  |  | C=O | 2877 |
|  |  | N-H | 1080 |
| 3 | Co-crystals | O-H stretching | 3749.74 |
|  |  | Other | 1458 [1489 for pure curcumin]<br>1026[1080for pure L-Proline] |
|  |  |  | 1080[1296 for pure Curcumin] |

The vibrational mode at 3410 cm−1, corresponding to the phenolic groups in the co-crystals, exhibited a minor displacement in comparison to pure curcumin and piperine [3495 and 3433 cm−1, respectively]. This shift can be attributed to pi-pi stacking interactions, signifying intermolecular connections between curcumin and piperine. In contrast, the alterations in the aromatic stretching peaks of curcumin within the co-crystals were minimal, indicating curcumin’s presence within the cocrystal structure. Further details are provided in **Table 4 and Figure 3B**.

**Table 4.** Observations of Fourier Transform-infrared spectra of (A) Curcumin(B) L-Piperine, and (C) the co-crystal of Curcumin-Piperine.

| Sr. No. | Sample | Bond | Wavenumber (cm-1) |
| --- | --- | --- | --- |
| 1 | Curcumin | Phenolic OH stretching | 3603 |
|  |  | Aromatic (C=O and C=C) stretching | 1509 and 1628 |
|  |  | C–H stretching | 1489 |
|  |  | Aromatic C-O | 1296 |
|  |  | C-O-C stretching | 949 |
| 2 | Piperine | Phenolic OH stretching | 3433 |
|  |  | Aromatic C-H stretching | 3178.79 |
|  |  | Aliphatic C-H stretching | 2985.91 |
|  |  | Carbonyl group (CO–N) | 1473.66 |
|  |  | Carbonyl C=O | 1666.55 |
| 3 | Co-crystals | Phenolic O–H stretching | 3410 cm<sup>–1</sup> [3495 and 3433 for pure Curcumin and piperine] |
|  |  | Carbonyl C=O | 1507 cm<sup>–1</sup> [1509 cm<sup>–1</sup> for pure Curcumin] |
|  |  | Carbonyl group (CO–N) | 1442.80 cm<sup>–1</sup> [1473.66 cm<sup>–1</sup> for pure Piperine] |

### 2.9. Differential Scanning Calorimetry of Curcumin-L-proline co-crystals

Pure curcumin was characterized, alongside co-former L-proline, and co-crystals using DSC. The pure drug exhibited a distinct endothermic peak at 178.02 °C, while L-proline displayed one at 225.62°C, aligning with their respective melting points. The co-crystal exhibited a melting point of 179.02 °C. Notably, the DSC analysis of the co-crystal did not reveal a peak corresponding to the melting of the co-former, confirming the co-crystal’s formation and the absence of a physical mixture. The alterations in thermal properties served as evidence for the successful co-crystal formation, as depicted in **Figure 4A**.

**Figure 4.**
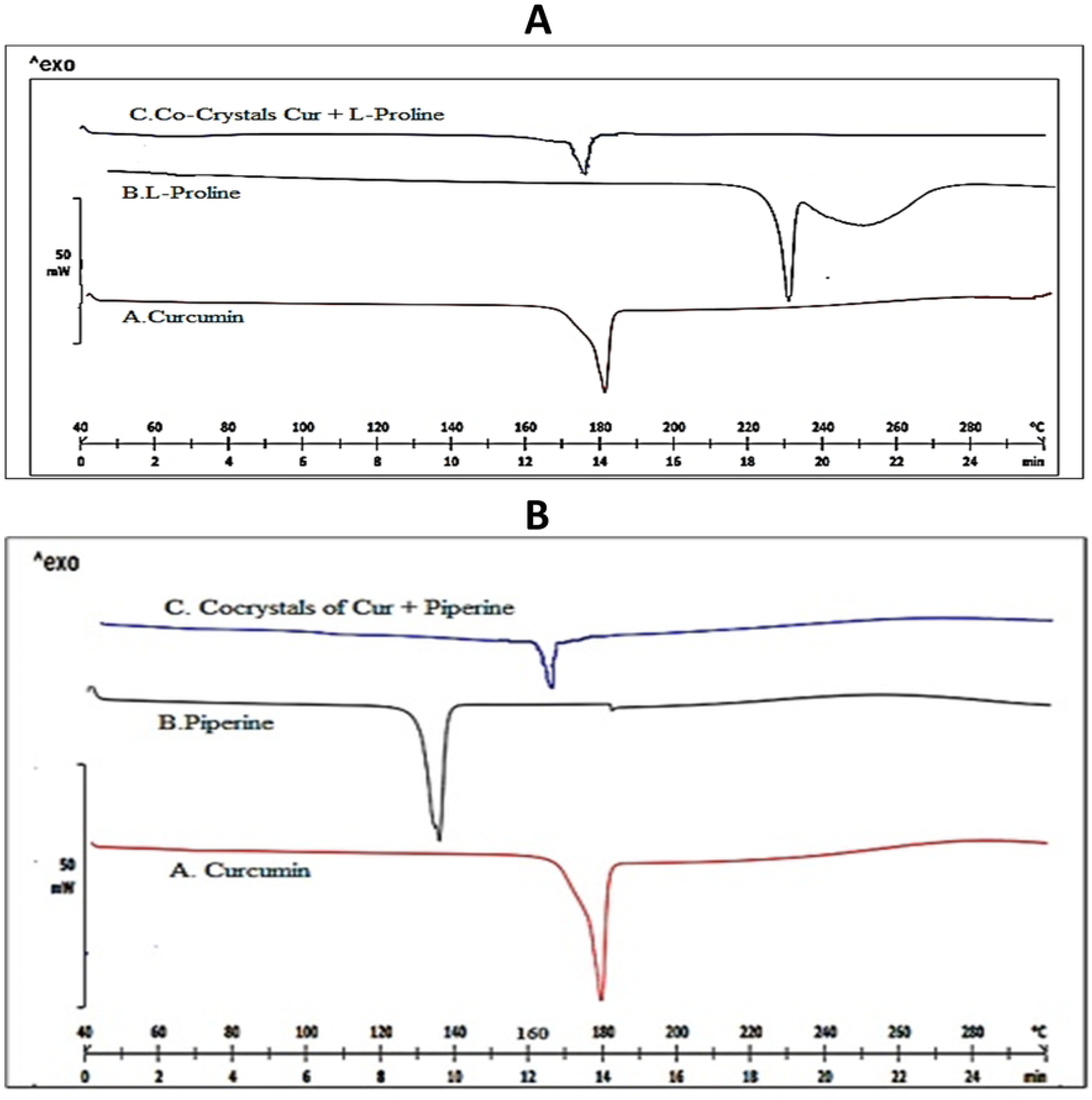
Differential Scanning Calorimetrythermogram of Curcumin, L-proline, and their respective co-crystals: (A) Curcumin-L-proline and (B) Curcumin-Piperine

Similarly, pure curcumin, co-former piperine, and cocrystals underwent DSC characterization. The pure drug demonstrated a characteristic endothermic peak at 178.02 °C, while the co-former piperine exhibited one at 133.65°C, consistent with their respective melting points. Remarkably, the cocrystal displayed a considerable deviation in melting point (163.20 °C) compared to the pure drug (178.02°C) and the co-former (133.65°C), as illustrated in **Figure 4B**.

### 2.10. Scanning Electron Microscopy (SEM)

The morphology of various substances including pure curcumin, pure L-proline, pure piperine, and co-crystals of curcumin – L-proline, as well as curcumin - piperine, is illustrated in **Figure 5**. Pure curcumin displays an irregular morphology, with sizes ranging from 4 to 15 µm. Similarly, both pure L-proline and piperine exhibit irregular morphologies, with an average size ranging between 1 to 4 µm. On the other hand, the co-crystals of curcumin – L-proline exhibit distinguishable shapes, which can be categorized into two types: elongated flakes and large spherical structures with some aggregates. The curcumin - L-proline co-crystals demonstrate a more pronounced distinction. They adopt a rod-like shape, accompanied by observed aggregates, within a size range of 2 µm – 5 µm.

**Figure 5.**
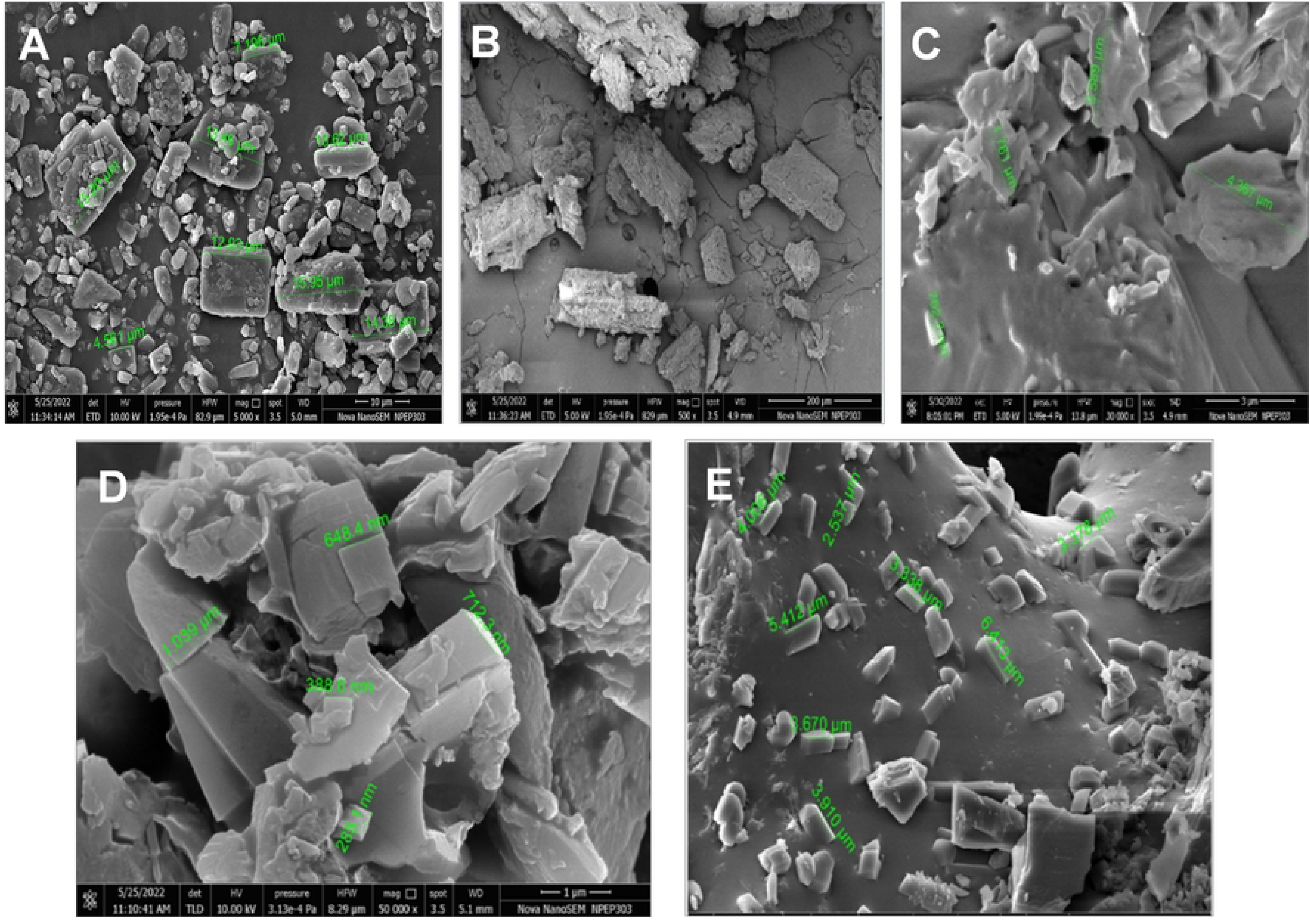
SEM images of (A) Curcumin (B) L-proline, (C) Piperine, (D) Co-crystals of curcumin – L-proline and (E) Co-crystals of curcumin –piperine

### 2.11. Powder X-ray diffraction Pattern

#### 2.11.1. PXRD of Curcumin and L-proline co-crystals

**Figure 6A** illustrates the PXRD patterns corresponding to curcumin, L-proline, and the co-crystal. These materials, while in a powdered state, exhibit distinct peaks of varying intensities at specific positions. The diffractogram of curcumin displays characteristic diffraction peaks at various 2θ values, indicative of its crystalline nature. Furthermore, **Table 5** presents the diffraction peaks obtained for both curcumin and the co-crystal with L-proline.

**Figure 6.**
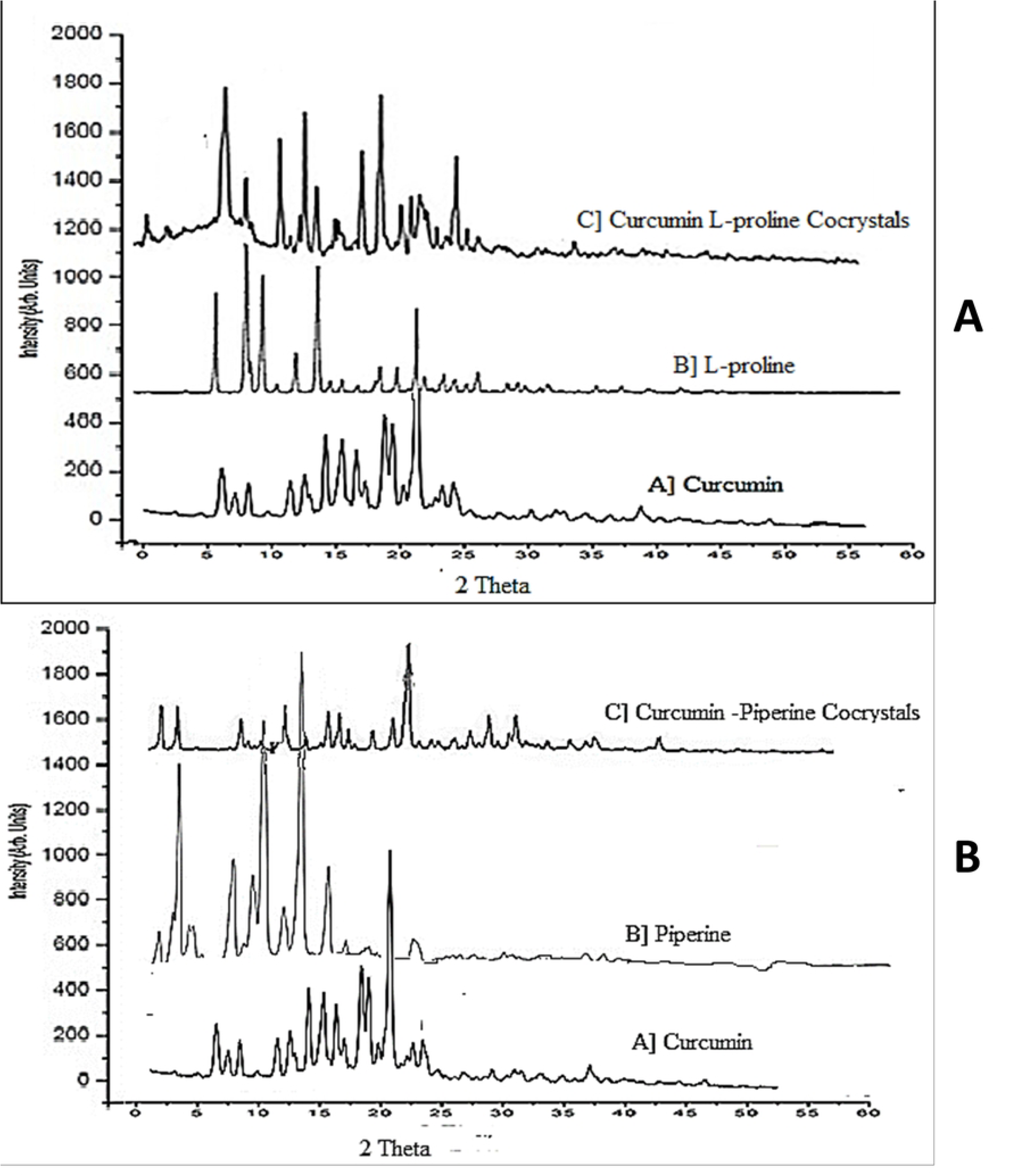
Powder X-ray diffraction Pattern of of Curcumin, L-proline, and their respective co-crystals: (A) Curcumin-L-proline and (B) Curcumin-Piperine

**Table 5.** Powder X-ray Diffraction peaks for Curcumin and L-Proline Cocrystals.

| Sr. No. | Compound | $2\theta$ value |
| --- | --- | --- |
| 1 | Curcumin | 7,8.9,12.2,14.5,17.4,18.2,21.2,23.4,23.9,24.6,25.6,26.1,26.8,27.3 and 29.0 |
| 2 | L-proline | 5.81, 11.57, 13.55, 14.20, 16.78, 26.35, 28.14 and 29.23 |
| 3 | Curcumin-L-proline co-crystals | 7.10, 12.28, 15.06, 18.28, 20.54, 21.67, 23.56 and 25.03. |

The PXRD pattern of the co-crystal stands apart from those of its individual components. Notably, additional diffraction peaks emerge, absent in the pure drug and co-former. **Table 6** outlines these extra diffraction peaks for the co-crystal at specific 2θ values. The presence of novel diffraction peaks within the co-crystal’s diffractogram signifies the formation of a new crystalline phase. Extensive documentation demonstrates the identification of co-crystals based on PXRD patterns, revealing distinct peaks that differ from those associated with its constituent compounds.

**Table 6.**
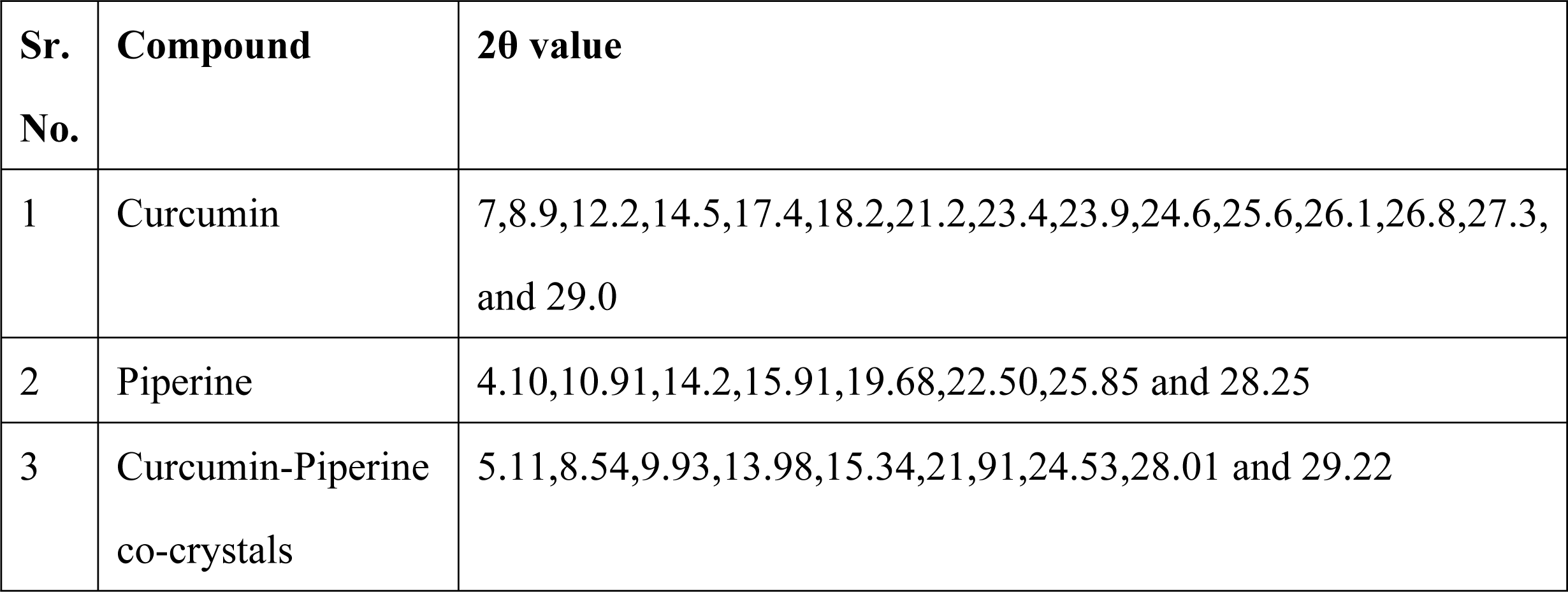
Powder X-ray Diffraction peaks for Curcumin and Piperine Cocrystals.

| Sr. No. | Compound | 2θ value |
| --- | --- | --- |
| 1 | Curcumin | 7,8.9,12.2,14.5,17.4,18.2,21.2,23.4,23.9,24.6,25.6,26.1,26.8,27.3, and 29.0 |
| 2 | Piperine | 4.10,10.91,14.2,15.91,19.68,22.50,25.85 and 28.25 |
| 3 | Curcumin-Piperine co-crystals | 5.11,8.54,9.93,13.98,15.34,21.91,24.53,28.01 and 29.22 |

#### 2.11.2. PXRD of Curcumin and Piperine co-crystals

**Figure 6B** illustrates the PXRD patterns corresponding to Curcumin, Piperine, and their co-crystals. The powdered materials exhibited distinct peaks with varying intensities at specific positions. The diffractogram of curcumin displayed characteristic diffraction peaks at various 2θ values, confirming its crystalline nature. Similarly, Piperine also exhibited identifiable diffraction peaks. Notably, the PXRD pattern of the co-crystal exhibited discernible differences from those of its individual components, revealing additional diffraction peaks absent in the pure drug and co-former. These supplementary diffraction peaks specific to the co-crystal were located at 2θ values as outlined in **Table 6**. The emergence of novel diffraction peaks in the co-crystal’s diffractogram signifies the creation of a new crystalline phase, namely the co-crystal. The formation of co-crystals, as evidenced by the PXRD pattern, has been extensively documented. These patterns demonstrate new peaks distinct from those corresponding to the constituent components, confirming the successful generation of co-crystals.

### 2.12. Evaluation of Formulation

#### 2.12.1. Pre-Compression Evaluation

The preformulation properties including angle of repose, bulk density, tapped density, Hauser’s ratio, and Carr’s index were determined to meet the acceptable criteria outlined in the USP. With Carr’s index near to 16% the powder shows reasonable compressibility and thus tablets were directly compressible.

#### 2.12.2. Post Compression Evaluation

The tablets that were formulated using cocrystals of curcumin met the standards outlined in the pharmacopoeia, as indicated in **Table 7**.

**Table 7.** Post Compression evaluation of immediate release tablets.

| Sr. No. | Parameter | Results |
| --- | --- | --- |
| 1 | Thickness (mm) | 2.09 ± 0.08 mm |
| 2 | Hardness (kg/cm<sup>2</sup>) | 4.5 (kg/cm<sup>2</sup>) |
| 3 | Friability (%) | 0.16% |
| 4 | Weight variation (%) | Less than 5% |
| 5 | Disintegration time | 3 min 15 sec |

#### 2.12.3. In-vitro dissolution study immediate release tablet

*In vitro* drug release results demonstrated an improved dissolution profile for the curcumin co-crystals IR tablet compared to the pure drug, as shown in Figure 7A. After 120 minutes, the curcumin co-crystals exhibited a drug release percentage of 86.56%, whereas the pure curcumin tablet only achieved a 24% drug release.

**Figure 7.**
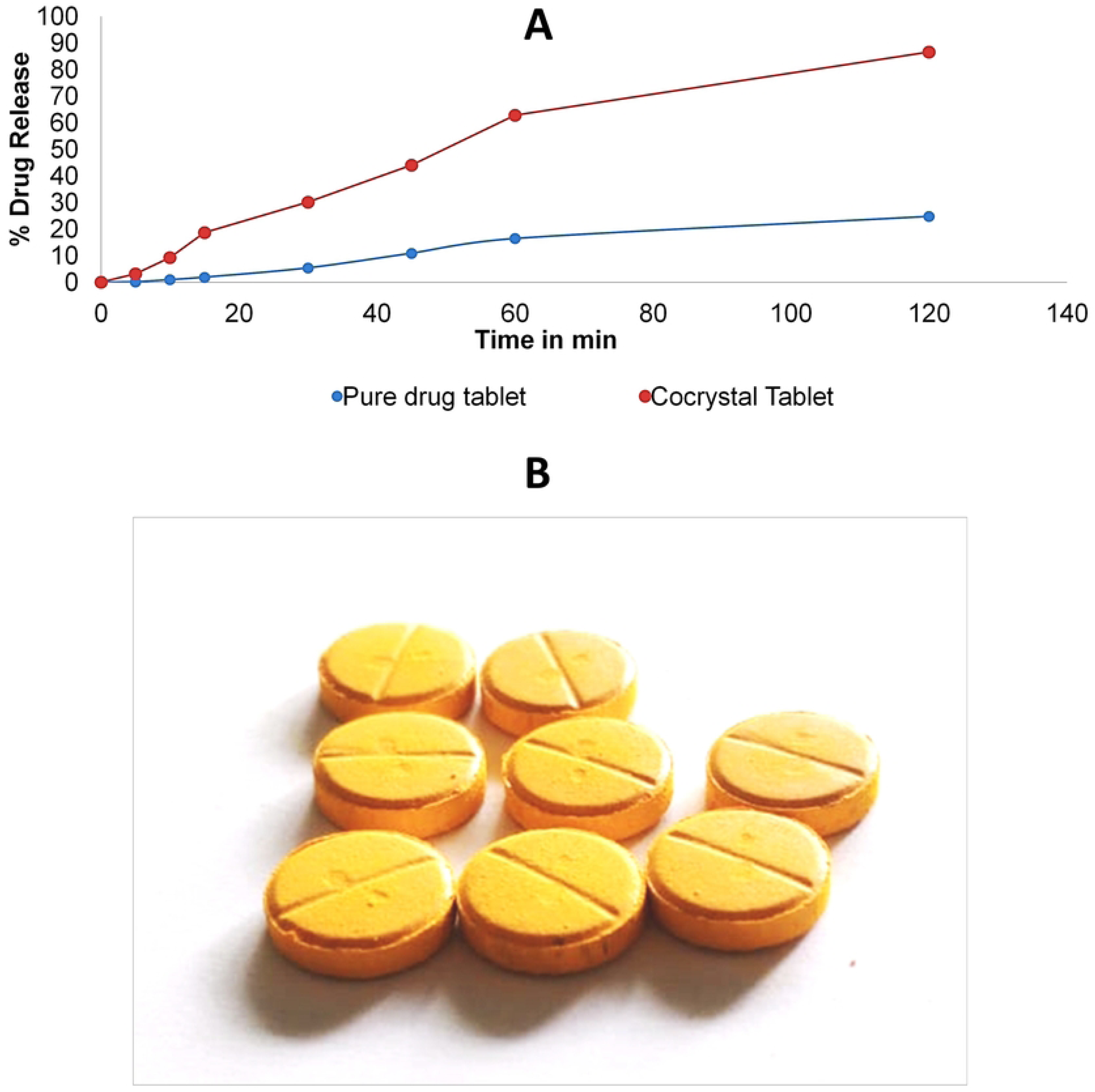
(A) In-vitro Drug Release Comparison between Immediate Release Pure Curcumin Tablets and Curcumin Co-crystal Tablets (B) Immediate Release Curcumin Tablets Incorporating Co-crystals.

#### 2.12.4. Stability Study

Tablets containing both co-crystals and individual co-crystals underwent a stability study under accelerated conditions for a duration of three months, aimed at evaluating their stability. The co-crystals exhibited stability under the designated storage conditions, and there were no significant alterations observed in key parameters such as melting point, solubility, and *in vitro* drug release after the three-month period at 25°C/60% RH, as presented in **Table 8**.

**Table 8.** Stability study data for tablets containing Co-crystals with L-proline at accelerated conditions for one month.

| Sr. No. | Parameter | Results | After 1 month |
| --- | --- | --- | --- |
| 1 | Colour & Appearance of tablets | Yellow | Yellow |
| 2 | In-vitro dissolution (%) from tablet | 86.56% | 86.5% |
| 3 | Drug content of tablets | 96.25 | 96.18 |

## 3. Materials and Methods

### 3.1. *In-silico* screening of the co-formers using molecular docking

Various tools and softwares like MGLTools1.5.6.,Marvin Sketch, Auto Dock 4.2.3 for docking process,PyRx,Open babel 2.3.2/GUI., Biovia discovery studio 2021 client version tool were used for in –silico screening of various coformers with curcumin.

The 3D structures of curcumin and all co-formers were acquired from Pub Chem (http://pubchem.ncbi.nih.gov) in the SDF (Structure Data Files) format. Subsequently, all SDF files were transformed into Mol2 format using Chem Draw software. The Mol2 files were then converted to PDB (Protein Data Bank) format using Auto Dock software. Afterwards, the PDB files were imported into Auto Dock 4.2.3, where they underwent conversion into pdbq files. This conversion included the addition of polar hydrogen atoms and Kollman charges. Following this, the pdbq files were further transformed into pdbqt format by calculating their torsion angles. These pdbqt files were then prepared for docking simulations. Lastly, the docking results were analyzed using both Auto Dock software and Discovery Studio.

### 3.2. Preparation of co-crystals

Curcumin and co-formers were combined in molar ratios of 1:1, 1:2, and 2:1. To dissolve the drug and co-former, 0.5 ml of methanol was introduced to the aforementioned mixtures. After thorough mixing and stirring, the solutions were covered with perforated aluminum foil to facilitate solvent evaporation. Subsequently, the mixtures were left undisturbed in a clean and dry location at room temperature for a duration of 2 days. Once dried, the resulting product was collected and stored in a desiccator until subsequent evaluation.

### 3.3. Optimization and characterization [28–31]

#### 3.3.1. Optimization of co-crystals using DoE

The experimental setup utilized Design Expert Software, version 13, incorporating a Central Composite Face-Centered Design (CCF) approach. The study considered two independent variables: the concentration of the drug (A) and the concentration of the co-former (B). A Central Composite Face-Centered Design (CCF) with a 2-factor, 3-level configuration was employed, setting the Alpha value to 1. In total, nine experimental runs were executed, encompassing three variables coded at three levels: high (+1), medium (0), and low (−1), that is 150,100 and 50 mg of curcumin and 46.5,31.2 and 15.6 mg of L proline and 115.5, 31.2 and 15.6 of piperine were the levels chosen. Distinct designs were chosen for curcumin-L-proline and curcumin-piperine batches, with the responses measured being the percentage release of the drug and the permeability in concentration mg/mL.The DOE suggested 9 batches for curcumin –Lproline as well as curcumin-piperine as shown in **Table 9** and **10**

**Table 9.** Suggested Nine batches by Designs of Experiment Software for solubility response.

| Standard | Run | Factor levels |  | Response |
| --- | --- | --- | --- | --- |
|  |  | Curcumin<br>(mg) | L-Proline<br>(mg) | % Drug Release |
| 2 | 1 | 150 | 15.6 | 25 |
| 5 | 2 | 50 | 31.2 | 30 |
| 10 | 3 | 100 | 31.2 | 71 |
| 1 | 4 | 50 | 15.6 | 40 |
| 6 | 5 | 150 | 31.2 | 43.6 |
| 4 | 6 | 150 | 46.8 | 50 |
| 8 | 7 | 100 | 46.8 | 30.6 |
| 7 | 8 | 100 | 15.6 | 22 |
| 3 | 9 | 50 | 46.8 | 25 |

**Table 10.** Suggested Nine batches by Designs of Experiment Software for Permeability response.

| Standard | Run | Factor levels |  | Response |
| --- | --- | --- | --- | --- |
|  |  | Curcumin<br>(mg) | Piperine<br>(mg) | Drug Permeated<br>mg/mL |
| 8 | 1 | 100 | 115.5 | 0.0401 |
| 6 | 2 | 150 | 77 | 0.0427 |
| 4 | 3 | 150 | 115.5 | 0.2034 |
| 5 | 4 | 100 | 77 | 0.2545 |
| 12 | 5 | 50 | 77 | 0.1030 |
| 2 | 6 | 150 | 38.5 | 0.0393 |
| 1 | 7 | 50 | 38.5 | 0.2036 |
| 3 | 8 | 50 | 115.5 | 0.0286 |
| 7 | 9 | 100 | 38.5 | 0.0303 |

#### 3.3.2. *In Vitro* dissolution rate profiles of cocrystals of curcumin -L-proline

The method of *in vitro* dissolution rate profiles reported earlier was adopted [32–35]. The in vitro dissolution profiles of both curcumin and co-crystals (for 9 batches as suggested by Design of Experiments) were assessed using a Type II paddle (USP) dissolution test apparatus. The apparatus was set to a rotation speed of 50 rpm in 900 mL of distilled water, which served as the dissolution medium. The temperature was carefully maintained at 37°C ± 0.5°C. Sampling was performed at specific time intervals (15, 30, 60, and up to 480 minutes / 8 hours). A 1 ml sample was withdrawn and then diluted with distilled water to a final volume of 10 ml.

The concentration of dissolved curcumin was determined by measuring the absorbance at the maximum wavelength of 425 nm using a UV-visible spectrophotometer. The collected data is presented in the form of a graph (time vs. % cumulative drug release) comparing the co-crystals’ % cumulative drug release with that of only curcumin.

#### 3.3.3. *Ex-vivo* permeability study for cocrystals of curcumin – piperine

Everted gut sac method was used to find out permeability of curcumin in cocrystal form with piperine. The jejunum part of the small intestine from a goat was identified and isolated. This segment was then placed in a petri dish containing phosphate buffer with a pH of 6.8 for thorough cleaning. After cleaning, the intestine was carefully turned inside out using a glass rod. Subsequently, a 3 cm piece of the jejunum section from the everted intestine was cut. One end of the piece was secured with a thread, and a dissected butterfly needle was inserted and tied at the other end, creating a gut sac. A solution with a pH of 6.8, totaling 2 ml, was introduced into the gut sac through the inserted butterfly needle. The everted intestinal sac was then placed in a beaker containing 100 ml of the experimental test solution and incubated.

#### 3.3.4. Saturation solubility Study

25 ml of distilled water (DW) was taken; to this, curcumin was added while continuously stirring until no more of the drug could dissolve. A slight excess of the drug was subsequently introduced. The flask was then placed on a magnetic stirrer for 24 hours and covered with aluminum foil to prevent light exposure. After the 24-hour stirring period, any lost water due to evaporation was compensated for by adjusting the volume, and the solution was filtered using Whatman filter paper. A 1 ml sample of the solution was further diluted with 10 ml of distilled water. The absorbance of the solution was measured at 425 nm using a UV-visible spectrophotometer.

#### 3.3.5. Fourier Transform-Infrared Spectroscopy Analysis

The intermolecular interactions of curcumin and co-formers were investigated both individually and in co-crystalline forms (curcumin & L-proline, curcumin & piperine) utilizing an FT-IR spectrophotometer. The samples were blended with dry potassium bromide, maintaining a weight ratio of 1:100. Subsequently, this mixture was compacted into pellets, and the infrared absorption spectra of the samples were measured in the range of 4000 cm−1 to 600 cm−1 [36–38].

#### 3.3.6. Differential Scanning Calorimetry (DSC)

Differential Scanning Calorimetry (DSC) was employed to document the melting behaviors of curcumin, co-formers, and co-crystals. A solid sample weighing 3-4 mg was carefully positioned on an aluminum pan. The pan was subsequently sealed with a lid and crimped securely using a DSC crimper. These crimp-sealed pans containing the solid samples were subjected to heating, ranging from 40°C to 280°C, with a heating rate of 10°C per minute. This process was carried out under a nitrogen flow of 17 ml/min, and the resulting endothermic peaks were analyzed.

#### 3.3.7. Scanning Electron Microscopy [SEM]

Scanning electron microscopy was employed to assess Curcumin, L-proline, piperine, as well as the co-crystals of Curcumin with L-proline and curcumin with piperine. The resulting images were utilized for both particle size determination and morphology analysis [39].

### 3.4. Formulation of Immediate release tablets

The final formulation for the curcumin co-crystals was selected to be immediate-release tablets **(Figure 7B)**. The formulation provided in **Table 11** was utilized, and the tablets were manufactured using the direct compression method. Compression was carried out using a 16- station rotary tablet compression machine equipped with 11mm round, flat punches.

**Table 11.** Composition of Curcumin Immediate release tablets containing Co-crystals and only curcumin.

| Sr. No. | Ingredients | Category | Quantity (mg) co-crystals of curcumin | Quantity (mg) curcumin |
| --- | --- | --- | --- | --- |
| 1 | Curcumin | Drug | 308.2 . Equivalent to [200mg Curcumin] | 200 |
| 2 | Micro crystalline cellulose | Diluent | 80 | 188.2 |
| 3 | Lactose monohydrate | Diluent | 61.8 | 61.8 |
| 4 | Sodium starch glycolate | Disintegrant | 40 | 40 |
| 5 | Magnesium stearate | Lubricant | 5 | 5 |
| 6 | Aerosil 200 | Glidant | 5 | 5 |
|  |  | Total | 500 | 500 |

Various pre-compression parameters, including the Angle of Repose, Bulk Density, Tapped Density, Compressibility Index, and Hausner’s Ratio, were assessed. Additionally, post-compression parameters such as Appearance, Thickness, Hardness, Weight Variation, Disintegration Time, *In-vitro* Dissolution Study, and Stability Study of the immediate-release tablets were conducted.

## 4. Conclusions

Curcumin, a promising phytochemical, faces limitations in practical applications due to its classification in BCS class IV, characterized by low water solubility and permeability. Enhancing the bioavailability of BCS class IV drugs is a significant challenge, but crystal chemistry offers a hopeful solution. In this study, Curcumin co-crystals were developed to improve both solubility and permeability. In contrast to traditional methods that involve labor-intensive experimentation and lengthy assessments of potential co-formers, molecular docking for *In-silico* co-former screening provides a systematic and logical approach to identifying suitable partners. Within this investigation, two unique co-crystals were created through solvent evaporation using methanol as the solvent at a 1:1 molar ratio. L-proline and piperine were selected as co-formers to enhance solubility and permeability. The application of *In-silico* screening through molecular docking proved highly effective and efficient for co-former selection. Specifically, the molecular ratios of Curcumin and L-proline (1:1) for solubility enhancement and Curcumin and piperine (1:1) for permeability improvement yielded promising results, supported by optimization analysis. After 8 hours, pure Curcumin exhibited a 20% cumulative drug release (CDR), while Curcumin-L-proline co-crystals achieved a remarkable 71% CDR. Additionally, the permeability of pure Curcumin was 0.0142 mg/mL, whereas the Curcumin-piperine combination showed increased permeability of 0.2545 mg/mL in a phosphate buffer at pH 6.8 after 4 hours. Formulating Curcumin co-crystals into immediate-release tablets resulted in an 86.56% drug release, compared to only 24% from tablets containing pure Curcumin. Furthermore, stability studies conducted according to ICH guidelines (40 ± 2°C and 75 ± 5% RH) indicated that the tablet formulation with Curcumin co-crystals remained stable for 3 months. These findings, supported by SEM, FTIR, DSC, Invitro dissolution and permeation study, underline the effective and scientific nature of the *In-silico* approach in co-former selection to enhance solubility, permeability, and stability of drugs across various BCS classes (II, III, and IV).

## 6. Declaration

### Data availability statement

All the data generated during the experiment are provided in the manuscript/supplementary material.

### Ethics approval statement

Not Applicable

### Patient consent statement

Not Applicable

### Permission to reproduce material from other sources

Not Applicable

### Clinical trial registration

Not Applicable

### Consent for participation/publication

Not Applicable

## Acknowledgements

Authors are thankful to the International Foundation for Collaborative Research (IFCR), Bangladesh for collaborative support.

## References

1. Medarević, D.P.; Kachrimanis, K.; Mitrić, M.; Djuriš, J.; Djurić, Z.; Ibrić, S. Dissolution Rate Enhancement and Physicochemical Characterization of Carbamazepine-Poloxamer Solid Dispersions. Pharmaceutical Development and Technology 2016, 21, 268–276, doi:10.3109/10837450.2014.996899.

2. Lipinski, C.A.; Lombardo, F.; Dominy, B.W.; Feeney, P.J. Experimental and Computational Approaches to Estimate Solubility and Permeability in Drug Discovery and Development Settings 1PII of Original Article: S0169-409X(96)00423-1. The Article Was Originally Published in Advanced Drug Delivery Reviews 23 (1997) 3–25. 1. *Advanced Drug Delivery Reviews* 2001, *46*, 3–26, doi:10.1016/S0169-409X(00)00129-0.

3. Lara-Ochoa, F.; Espinosa-Pérez, G. Cocrystals Definitions. Supramolecular Chemistry 2007, 19, 553–557, doi:10.1080/10610270701501652.

4. Serajuddin, A.T.M. Salt Formation to Improve Drug Solubility. Advanced Drug Delivery Reviews 2007, 59, 603–616, doi:10.1016/j.addr.2007.05.010.

5. Bighley LD, Berge SM, Monkhouse DC. Encyclopedia of Pharmaceutical Technology. Vol. 1996;

6. Porter, C.J.H.; Trevaskis, N.L.; Charman, W.N. Lipids and Lipid-Based Formulations: Optimizing the Oral Delivery of Lipophilic Drugs. Nat Rev Drug Discov 2007, 6, 231–248, doi:10.1038/nrd2197.

7. Gadade, D.D.; Pekamwar, S.S. Pharmaceutical Cocrystals: Regulatory and Strategic Aspects, Design and Development. Adv Pharm Bull 2016, 6, 479–494, doi:10.15171/apb.2016.062.

8. Kumari, N.; Ghosh, A. Cocrystallization: Cutting Edge Tool for Physicochemical Modulation of Active Pharmaceutical Ingredients. CPD 2020, 26, 4858–4882, doi:10.2174/1381612826666200720114638.

9. Roy, P.; Ghosh, A. Mechanochemical Cocrystallization to Improve the Physicochemical Properties of Chlorzoxazone. CrystEngComm 2020, 22, 4611–4620, doi:10.1039/D0CE00635A.

10. Kumar, S.; Nanda, A. Pharmaceutical Cocrystals: An Overview. pharmaceutical-sciences 2017, 79, doi:10.4172/pharmaceutical-sciences.1000302.

11. Alt, A.; Potthast, H.; Moessinger, J.; Sickmüller, B.; Oeser, H. Biopharmaceutical Characterization of Sotalol-Containing Oral Immediate Release Drug Products. European Journal of Pharmaceutics and Biopharmaceutics 2004, 58, 145–150, doi:10.1016/j.ejpb.2004.02.007.

12. Good, D.J.; Rodríguez-Hornedo, N. Solubility Advantage of Pharmaceutical Cocrystals. Crystal Growth & Design 2009, 9, 2252–2264, doi:10.1021/cg801039j.

13. Karimi-Jafari, M.; Padrela, L.; Walker, G.M.; Croker, D.M. Creating Cocrystals: A Review of Pharmaceutical Cocrystal Preparation Routes and Applications. Crystal Growth & Design 2018, 18, 6370–6387, doi:10.1021/acs.cgd.8b00933.

14. Song, Y.; Wang, L.-Y.; Liu, F.; Li, Y.-T.; Wu, Z.-Y.; Yan, C.-W. Simultaneously Enhancing the *in Vitro* / *in Vivo* Performances of Acetazolamide Using Proline as a Zwitterionic Coformer for Cocrystallization. CrystEngComm 2019, 21, 3064–3073, doi:10.1039/C9CE00270G.

15. Yan, Y.; Chen, J.-M.; Lu, T.-B. Simultaneously Enhancing the Solubility and Permeability of Acyclovir by Crystal Engineering Approach. CrystEngComm 2013, 15, 6457, doi:10.1039/c3ce41017j.

16. Bolla, G.; Nangia, A. Pharmaceutical Cocrystals: Walking the Talk. Chem. Commun. 2016, 52, 8342–8360, doi:10.1039/C6CC02943D.

17. Roy, P.; Ghosh, A. Progress on Cocrystallization of Poorly Soluble NME’s in the Last Decade. CrystEngComm 2020, 22, 6958–6974, doi:10.1039/D0CE01276A.

18. Alhalaweh, A.; Roy, L.; Rodríguez-Hornedo, N.; Velaga, S.P. PH-Dependent Solubility of Indomethacin–Saccharin and Carbamazepine–Saccharin Cocrystals in Aqueous Media. *Mol*. Pharmaceutics 2012, 9, 2605–2612, doi:10.1021/mp300189b.

19. Bavishi, D.D.; Borkhataria, C.H. Spring and Parachute: How Cocrystals Enhance Solubility. Progress in Crystal Growth and Characterization of Materials 2016, 62, 1–8, doi:10.1016/j.pcrysgrow.2016.07.001.

20. Ghadi, R.; Dand, N. BCS Class IV Drugs: Highly Notorious Candidates for Formulation Development. Journal of Controlled Release 2017, 248, 71–95, doi:10.1016/j.jconrel.2017.01.014.

21. Thakuria, R.; Sarma, B. Drug-Drug and Drug-Nutraceutical Cocrystal/Salt as Alternative Medicine for Combination Therapy: A Crystal Engineering Approach. Crystals 2018, 8, 101, doi:10.3390/cryst8020101.

22. Modani, S.; Gunnam, A.; Yadav, B.; Nangia, A.K.; Shastri, N.R. Generation and Evaluation of Pharmacologically Relevant Drug–Drug Cocrystal for Gout Therapy. Crystal Growth & Design 2020, 20, 3577–3583, doi:10.1021/acs.cgd.0c00106.

23. Guo, M.; Sun, X.; Chen, J.; Cai, T. Pharmaceutical Cocrystals: A Review of Preparations, Physicochemical Properties and Applications. Acta Pharmaceutica Sinica B 2021, 11, 2537– 2564, doi:10.1016/j.apsb.2021.03.030.

24. Ghafari, S.; Aziz, H.A.; Isa, M.H.; Zinatizadeh, A.A. Application of Response Surface Methodology (RSM) to Optimize Coagulation–Flocculation Treatment of Leachate Using Poly-Aluminum Chloride (PAC) and Alum. Journal of Hazardous Materials 2009, 163, 650–656, doi:10.1016/j.jhazmat.2008.07.090.

25. Buddhadev, S.S.; Garala, K.C. Pharmaceutical Cocrystals—A Review. In Proceedings of the The 2nd International Online Conference on Crystals; MDPI, March 8 2021; p. 14.

26. Kumar, S.; Nanda, A. Pharmaceutical Cocrystals: An Overview. pharmaceutical-sciences 2017, 79, doi:10.4172/pharmaceutical-sciences.1000302.

27. Taylor, C.R.; Day, G.M. Evaluating the Energetic Driving Force for Cocrystal Formation. Crystal Growth & Design 2018, 18, 892–904, doi:10.1021/acs.cgd.7b01375.

28. Fukuda, I.M.; Pinto, C.F.F.; Moreira, C.D.S.; Saviano, A.M.; Lourenço, F.R. Design of Experiments (DoE) Applied to Pharmaceutical and Analytical Quality by Design (QbD). *Braz*. J. Pharm. Sci. 2018, 54, doi:10.1590/s2175-97902018000001006.

29. Gunst, R.F. Response Surface Methodology: Process and Product Optimization Using Designed Experiments. Technometrics 1996, 38, 284–286, doi:10.1080/00401706.1996.10484509.

30. Asghar, A.; Abdul Raman, A.A.; Daud, W.M.A.W. A Comparison of Central Composite Design and Taguchi Method for Optimizing Fenton Process. The Scientific World Journal 2014, 2014, 1–14, doi:10.1155/2014/869120.

31. Bashiri, M.; Farshbaf Geranmayeh, A. Tuning the Parameters of an Artificial Neural Network Using Central Composite Design and Genetic Algorithm. Scientia Iranica 2011, 18, 1600–1608, doi:10.1016/j.scient.2011.08.031.

32. Newa, M.; Bhandari, K.H.; Kim, J.O.; Im, J.S.; Kim, J.A.; Yoo, B.K.; Woo, J.S.; Choi, H.G.; Yong, C.S. Enhancement of Solubility, Dissolution and Bioavailability of Ibuprofen in Solid Dispersion Systems. Chem. Pharm. Bull. 2008, 56, 569–574, doi:10.1248/cpb.56.569.

33. Development of the NSAID-L-Proline Amino Acid Zwitterionic Cocrystals. japs 2018, 57– 63, doi:10.7324/JAPS.2018.8408.

34. Sanii, R.; Andaloussi, Y.H.; Patyk-Kaźmierczak, E.; Zaworotko, M.J. Polymorphism in Ionic Cocrystals Comprising Lithium Salts and L -Proline. Crystal Growth & Design 2022, 22, 3786–3794, doi:10.1021/acs.cgd.2c00172.

35. Nugrahani, I.; Auli, W.N. Diclofenac-Proline Nano-Co-Crystal Development, Characterization, in Vitro Dissolution and Diffusion Study. Heliyon 2020, 6, e04864, doi:10.1016/j.heliyon.2020.e04864.

36. Deshkar, S.; Satpute, A. FORMULATION AND OPTIMIZATION OF CURCUMIN SOLID DISPERSION PELLETS FOR IMPROVED SOLUBILITY. Int J App Pharm 2019, 36–46, doi:10.22159/ijap.2020v12i2.34846.

37. Nascimento, A.L.C.S.; Fernandes, R.P.; Charpentier, M.D.; Ter Horst, J.H.; Caires, F.J.; Chorilli, M. Co-Crystals of Non-Steroidal Anti-Inflammatory Drugs (NSAIDs): Insight toward Formation, Methods, and Drug Enhancement. Particuology 2021, 58, 227–241, doi:10.1016/j.partic.2021.03.015.

38. Wünsche, S.; Seidel-Morgenstern, A.; Lorenz, H. Cocrystallization of Curcuminoids with Hydroxybenzenes Pyrogallol and Hydroxyquinol: Investigations of Binary Thermal Phase Behaviors. Crystal Growth & Design 2022, 22, 3303–3310, doi:10.1021/acs.cgd.2c00123.

39. Renza-Diaz, V.; Gonzalez-Hernández, M.; Pantoja, K.D.; D’Vries, R.F. Mechanochemical Treatment of Quercetin and Curcumin to Obtain Eutectic Mixtures with High Antioxidant Activity. CrystEngComm 2021, 23, 4985–4993, doi:10.1039/D1CE00697E.

